# A machine learning approach to quantify individual gait responses to ankle exoskeletons

**DOI:** 10.1101/2023.01.20.524757

**Authors:** Megan R. Ebers, Michael C. Rosenberg, J. Nathan Kutz, Katherine M. Steele

**Author notes:** (M.R. Ebers).

## Abstract

We currently lack a theoretical framework capable of characterizing heterogeneous responses to exoskeleton interventions. Predicting an individual’s response to an exoskeleton and understanding what data are needed to characterize responses has been a persistent challenge. In this study, we leverage a neural network-based discrepancy modeling framework to quantify complex changes in gait in response to passive ankle exoskeletons in nondisabled adults. Discrepancy modeling aims to resolve dynamical inconsistencies between model predictions and real-world measurements. Neural networks identified models of (i) *Nominal* gait, (ii) *Exoskeleton* (*Exo*) gait, and (iii) the *Discrepancy* (*i.e.*, response) between them. If an *Augmented* (Nominal+Discrepancy) model captured exoskeleton responses, its predictions should account for comparable amounts of variance in *Exo* gait data as the *Exo* model. Discrepancy modeling successfully quantified individuals’ exoskeleton responses without requiring knowledge about physiological structure or motor control: a model of *Nominal* gait augmented with a *Discrepancy* model of response accounted for significantly more variance in *Exo* gait (median *R*^2^ for kinematics (0.928 – 0.963) and electromyography (0.665 – 0.788), (*p* < 0.042)) than the *Nominal* model (median *R*^2^ for kinematics (0.863 – 0.939) and electromyography (0.516 – 0.664)). However, additional measurement modalities and/or improved resolution are needed to characterize *Exo* gait, as the discrepancy may not comprehensively capture response due to unexplained variance in *Exo* gait (median *R*^2^ for kinematics (0.954 – 0.977) and electromyography (0.724 – 0.815)). These techniques can be used to accelerate the discovery of individual-specific mechanisms driving exoskeleton responses, thus enabling personalized rehabilitation.

## 1. Introduction

Ankle exoskeletons and other assistive devices augment locomotion in nondisabled adults and improve kinematics, provide musculoskeletal alignment, and reduce the energetic demands of walking for individuals with neurologic injuries (Brehm et al., 2008; Maltais et al., 2001; Zhang et al., 2017; McCain et al., 2019; Conner et al., 2020; Collins et al., 2015; Ding et al., 2018; Koller et al., 2015; Malcolm et al., 2013). However, it is difficult to predict how exoskeleton forces and torques acting on the body alter an individual’s gait biomechanics, motor control, and sensory feedback (Steele et al., 2015; Jackson et al., 2017; Nuckols et al., 2020). Specifically, an incomplete understanding of the factors driving heterogeneous responses to exoskeletons makes identifying exoskeleton design parameters that optimize an individual’s gait challenging (McCain et al., 2019; Ries et al., 2015; Jackson and Collins, 2015; Kerkum et al., 2015; Collins et al., 2015). The most successful methods for personalizing exoskeletons are experimental techniques – such as human-in-the-loop optimization – that are resource intensive and iterative, placing additional burden on participants (Zhang et al., 2017; Ding et al., 2018; Ingraham et al., 2022). Conversely, computational approaches require shorter experimental sessions, but are currently unsuccessful due to the poor modeling of individual differences in neuromuscular physiology and control (Rosenberg and Steele, 2017; Lerner et al., 2019a; Afschrift et al., 2014; Crabtree and Higginson, 2009; Arch et al., 2016). Despite diverse approaches to characterizing exoskeleton impacts on gait, predicting responses to exoskeletons (*i.e.*, complex changes in gait with changes in exoskeleton properties) is an open problem (Kuo, 2002; Collins et al., 2005; Kuo et al., 2005; Uchida et al., 2016; Bregman et al., 2011; Collins et al., 2015; Pitto et al., 2019; Koller et al., 2015; Ries et al., 2015; Rosenberg and Steele, 2017; Rosenberg et al., 2020; Sawicki and Khan, 2015; Jackson and Collins, 2015; Jackson et al., 2017; Jacobs et al., 2018; Lerner et al., 2019b; Ding et al., 2018; Franks et al., 2020; Veerkamp et al., 2019; Meyer et al., 2017; Sauder et al., 2019). Emerging machine learning methods may help overcome experimental and theoretical barriers to quantifying individual-specific responses to exoskeletons.

While studies have investigated potential physiological mechanisms underlying exoskeleton responses, the complexity of the neuromusculoskeletal system limits our ability to uncover individual-specific physiological mechanisms governing exoskeleton gait (Sawicki and Khan, 2015; Jackson et al., 2017; Pitto et al., 2019; Falisse et al., 2019). For example, muscle-tendon mechanics are known to explain unexpected effects of exoskeleton assistance on gait energetics (Sawicki and Khan, 2015; Jackson et al., 2017). This mechanism alone, however, is unlikely to explain responses across individuals, which may be influenced by other factors, such as sensation or motor control (Yandell et al., 2017; Pitto et al., 2019). Other researchers have used physics-based or data-driven models to capture the physiological processing driving exoskeleton responses without explicit physiological mechanisms (Rosenberg et al., 2020; Ries et al., 2014, 2015). For example, a Random Forest algorithm with kinematic and clinical exam measurements from over 300 children with cerebral palsy was used to predict changes in kinematics with passive ankle exoskeletons (Ries et al., 2014). The algorithm, however, only explained 19-28% of the variance in kinematic responses. Quantifying individual-specific processes – complex interactions between biomechanical, neural, and sensory mechanisms – that can explain heterogeneous exoskeleton response could inform personalized device designs or improve accuracy of gait simulations with exoskeletons.

An unexplored approach to accelerate discovery of individual-specific processes driving complex gait changes with exoskeletons is characterizing the response itself. Gait kinematics, kinetics, and muscle activity change with exoskeletons, and our inability to explain this response represents a *discrepancy* in our understanding of the neuro-musculoskeletal system and how it responds to an exoskeleton. This discrepancy may reflect many individual factors, such as inter-individual differences in musculoskeletal physiology or motor control, an incomplete understanding of gait adaptation with exoskeletons, or an inability to model these complex processes. It is precisely because *we do not know* what processes (or combinations thereof) underlie responses that, in this paper, we choose to summarize an all-encompassing discrepancy to describe exoskeleton responses. Additionally, it is unknown what data are needed to capture discrepancies. In previous studies, some researchers used only a few strides to tune models (Meyer et al., 2017; Sauder et al., 2019; Pitto et al., 2019), while others used large datasets (Rosenberg et al., 2020). In tuning models, a variety of data were used: kinematics (Delp et al., 2007), electromyography (EMG) (Meyer et al., 2017; Serrancolí et al., 2016), and clinical exams (Pitto et al., 2019; Ries et al., 2014), among others. Quantifying this discrepancy is, therefore, a critical step in understanding individual responses to ankle exoskeletons.

Discrepancy modeling is a novel tool in machine learning developed to identify missing physics in complex systems described by differential equations (Ebers et al., 2022), such as those describing human movement (Hatze, 1977, 1976; Maus et al., 2015; Mert et al., 2015; Wang et al., 2016). A variety of machine learning techniques can be used to quantify discrepancies (Ebers et al., 2022). Deep learning is a sub-field of machine learning, with neural networks comprising the backbone of deep learning algorithms. We employ a simple neural network underpinning more complicated deep learning architectures (Goodfellow et al., 2016). Neural networks have been used to model complex behavior from time-series gait data (LeCun et al., 2015; Rosenberg et al., 2020; Winner et al., 2022), making them an ideal candidate to model exoskeleton responses as discrepancies. Neural networks (i) can learn complex patterns in experimental data that coalesce from diverse sources, (ii) do not make explicit assumptions about physiology and motor control, and (iii) learn time-variant features of exploration. In biomechanics research, neural networks have been used to classify and predict human movement (Sansano et al., 2020; Hernandez et al., 2021; Cao et al., 2017; Mathis et al., 2018; Baccouche et al., 2011). While neural networks applied to lower-limb exoskeletons and prostheses are most often used for trajectory prediction and device control (Liu et al., 2017; Zaroug et al., 2020; Su and Gutierrez-Farewik, 2020; Kolaghassi et al., 2022; Rosenberg et al., 2020; Vu et al., 2020), they have not quantified exoskeleton responses.

The purpose of this study was to determine if a discrepancy modeling framework could quantify individual-specific gait responses to ankle exoskeletons (Fig. 1). We employed neural network-based discrepancy modeling to encode processes governing joint kinematic and EMG responses of nondisabled adults to bilateral passive ankle exoskeletons. Specifically, we modeled the discrepancy between the processes governing gait in (i) a *Nominal Condition* (*i.e.*, zero-stiffness exoskeletons) and (ii) an *Exo Condition* (*i.e.*, exoskeletons with stiff springs resisting ankle dorsiflexion). If a *Discrepancy* model can encode exoskeleton responses, then augmenting a model of *Nominal* gait with the *Discrepancy* model should predict *Exo* kinematics and EMG more accurately than the *Nominal* model alone. Therefore, we hypothesized that (i) the *Nominal* model would predict *Exo* kinematics and EMG less accurately than for the *Nominal* condition, and (ii) the *Augmented* (Nominal+Discrepancy) model would capture greater variance in *Exo* kinematics and EMG than the *Nominal* model. To assess whether standard gait measurements have sufficient resolution for encoding exoskeleton gait, we evaluated the extent to which the *Exo* model could predict *Exo* gait. Finally, to assess the viability of discrepancy modeling in gait analysis settings, we evaluated the effect of limiting training data quantity on predictions of *Exo* gait.

**Figure 1:**
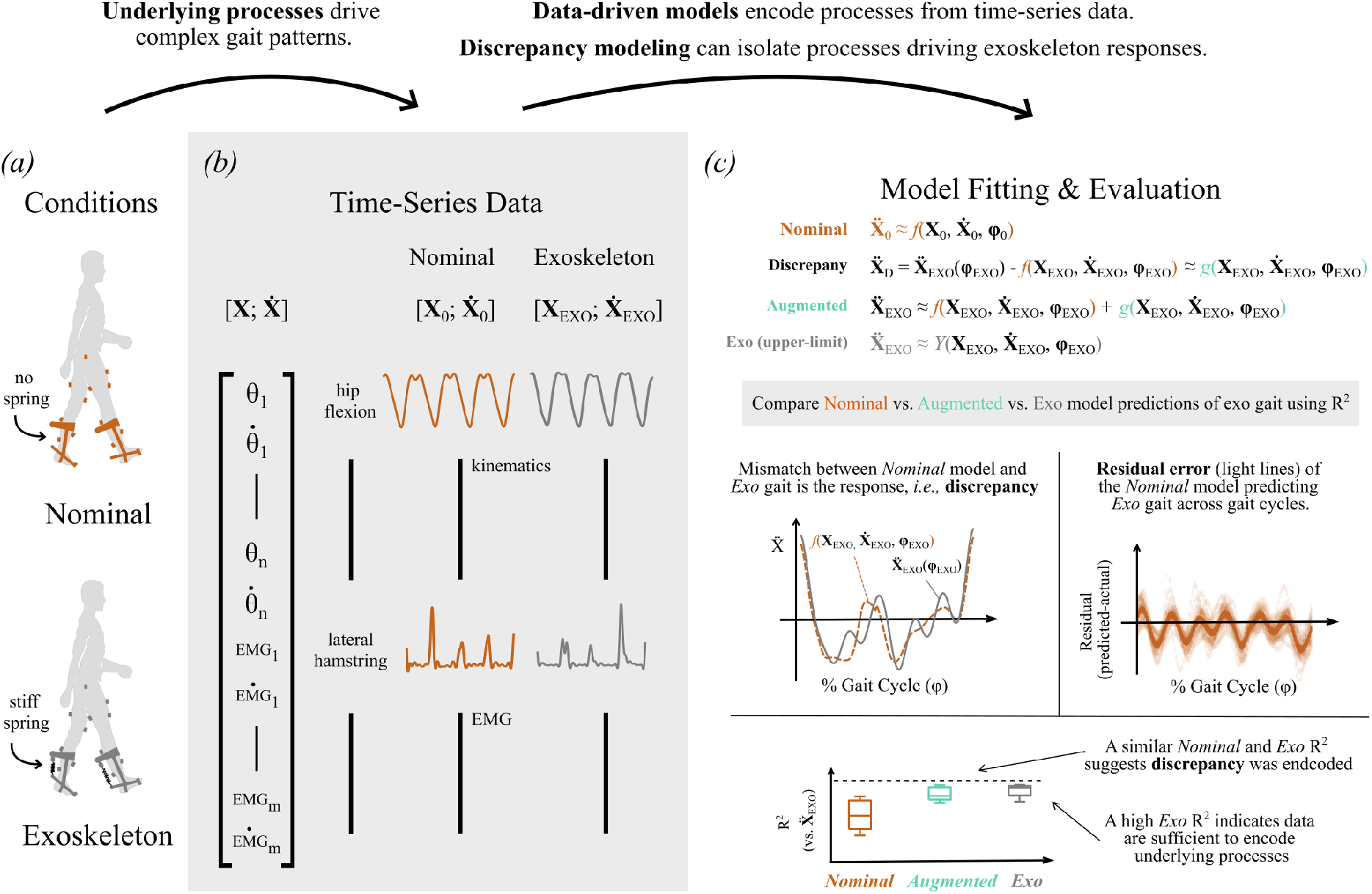
Framework outlining a machine learning approach using discrepancy modeling to quantify individual exoskeleton responses. *(a)* Joint kinematics (*θ*) and electromyography (EMG) data were collected from nondisabled participants during treadmill walking in bilateral passive ankle exoskeletons. Two conditions were analyzed: Nominal (exoskeleton with no spring) and *Exo* (exoskeleton with a 5 *N m deg*^-1^ spring). *(b)* We encoded the processes governing the time evolution of gait by identifying a data-driven differential equation function transforming model inputs (time-series gait data, **X**, their first time derivatives, 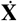, and percent stride, *ϕ*) into their outputs (second time derivatives of gait, 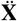). *(c)* Feed-forward neural network models identified: (1) a **Nominal model** (orange) using Nominal gait data, **X**_0_. The discrepancy between *Exo* gait and the *Nominal* model predictions is the exoskeleton response. (2) A **Discrepancy model** was trained using the identified response for each individual. This *Discrepancy* model was added to the *Nominal* model to create an **Augmented model** (green). (3) Finally, an **Exo model** (gray) was trained using the *Exo* gait data, **X**_*EXO*_; this served as the maximum expected variance. We compared how much variance in *Exo* gait is accounted for (*R*^2^) using the *Nominal, Augmented* (Nominal+Discrepancy), and *Exo* models and performed residual analysis on model predictions.

## 2. Methods

### 2.1. Human subjects

We analyzed gait data for 12 nondisabled adults (6F/6M; age = 23.9 ± 1.8 yrs; height = 1.69 ± 0.10 m; mass = 66.5 ± 11.7 kg) during treadmill walking in bilateral passive ankle exoskeletons at a self-selected speed (speed = 1.36 ± 0.11 m/s; data available: https://simtk.org/projects/ankleexopred) (Rosenberg et al., 2020). Exoskeletons resisted ankle dorsiflexion using linear springs attached in parallel to the shank. Data were captured in two conditions: The *Nominal Condition* involved walking while wearing exoskeletons without springs resisting ankle flexion (*i.e.*, zero stiffness); The *Exo Condition* involved walking in exoskeletons resisting ankle dorsiflexion with a stiffness of 5 *N m deg*^-1^ (Fig. 1a).

The complete experimental protocol, including exoskeleton fitting and walking practice, is described in (Rosenberg et al., 2020). Briefly, after an exoskeleton fitting and practice session on a prior day, participants walked on a split-belt instrumented treadmill (Bertec Corp., Columbus, USA) for six minutes per condition: a two-minute familiarization period, followed by four minutes at their self-selected speed. Conditions were randomized. Marker motion was recorded using a 10-camera optical motion capture system (Qualisys AB, Gothenburg, SE) and EMG signals were recorded bilaterally from five muscles (vastus medialis, soleus, gastrocnemius, gluteus medius, lateral hamstrings) (Delsys Inc., Natick, USA). Joint kinematics were estimated from marker data using the Inverse Kinematics algorithm in OpenSim 3.3 with a 19 degree-of-freedom skeletal model (Rajagopal et al., 2018; Hicks et al., 2015). Joint kinematics were low-pass filtered at 6Hz using a fourth-order Butterworth filter (Rosenberg et al., 2020). EMG data were high-pass filtered at 40Hz, rectified, and low-pass filtered at 10Hz using fourth-order Butterworth filters. All human subjects procedures were evaluated by the Institutional Review Board at the University of Washington.

### 2.2. Continuous-time neural network models of walking with ankle exoskeletons

To characterize discrepancies between processes underlying *Nominal* and *Exo* gait, we fit individual-specific neural networks to each participant’s data. The neural networks learned continuous-time nonlinear transformations from joint kinematics and EMG to their time derivatives, thereby encoding the biomechanical, neural, and sensory processes driving the time evolution of gait and how those processes change with ankle exoskeletons (Hartman, 2002; Guckenheimer and Holmes, 2013; Hartman, 1982). Input data for the *Nominal*, *Exo*, and *Discrepancy* models included the times series of 10 lower-limb joint angles, their angular velocities, 14 EMG signals, and their first time derivatives; percent stride, *ϕ*, was also included as an input (Fig. 1b). These variables reflect common measurements in gait analysis that are relevant to quantifying and synthesizing human movement (Collins et al., 2015; Kerkum et al., 2015; Zajac et al., 2002; Geyer and Herr, 2010; Rosenberg et al., 2020; Clark et al., 2010; Steele et al., 2017). Output data for the *Nominal* and *Exo* models included hip, knee, and ankle angular accelerations and the second time derivatives of the EMG signals from their respective exoskeleton condition. Output data for the *Discrepancy* model was the exoskeleton response, as identified in Eq. 3. We partitioned training and test sets by holding out the final 30 seconds (12.5%) of the data.

For each participant, we identified feed-forward neural network models (3 layers with activation functions logsig / radbas / purelin; 64 nodes per layer; MATLAB, Deep Learning Toolbox) (Fig. 1c) of walking for the Eq. 1: *Nominal Condition*, Eq. 2: *Exo Condition*, and Eq. 3: the *Discrepancy* between the *Nominal* and *Exo* conditions:

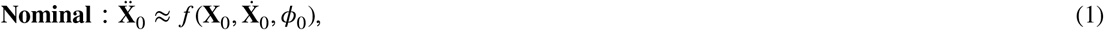

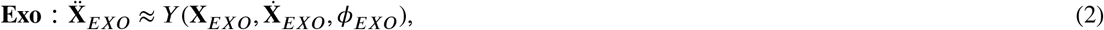

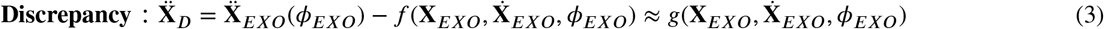

The *Discrepancy* model can then be used to *Augment* the *Nominal* model:

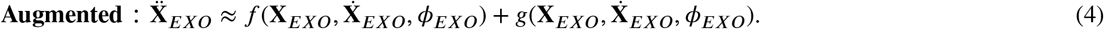

By adding the *Discrepancy* model to the *Nominal* model, we are testing whether there are organized and predictable features captured in the *Discrepancy* that can be used to quantify differences in individual exoskeleton responses. If the kinematic and EMG data are sufficient to capture responses, then the *Augmented* model (Eq. 4) should predict *Exo* gait as accurately as the *Exo* model (Eq. 2). The *Exo* model thus serves as an idealized ‘upper-limit’ case (*i.e.*, predictive potential when the response is known).

### 2.3. Statistical analysis

We quantified, using the coefficient of determination (*R*^2^), the extent to which processes underlying *Nominal, Augmented* (Nominal+Discrepancy), and *Exo* models could predict the held-out joint angle accelerations and second time derivatives of EMG during *Exo* gait. For each variable and across individuals, we compared *R*^2^ between (i) *Nominal* and *Augmented* models, (ii) *Nominal* and *Exo* models, and (iii) *Augmented* and *Exo* models using Wilcoxon Signed-Rank tests with Holm-Sidak Stepdown corrections for multiple comparisons (*α* = 0.05) (Rosenberg et al., 2020; Glantz, 2002). One participant was excluded from analysis due to changes in gait – possibly due to adaptation or conscious exploration of different gait patterns – late in the *Exo* condition walking trial. Additionally, we compared variance accounted for by the *Exo* model against *R*^2^ = 1. If these data alone could explain *Exo* gait, *R*^2^ would approach 1, and *R*^2^ < 1 would suggest additional measurement modalities or higher-resolution signals are needed to encode *Exo* gait.

Additionally, we performed residual analysis on model predictions (Cook and Weisberg, 1982; Belloto and Sokolovski, 1985). We plotted residuals across all gait cycles for the *Nominal*, *Augmented* (Nominal+Discrepancy), and *Exo* model predictions. Non-zero stride-averaged residuals indicate that a model does not capture processes driving gait. If the stride-averaged residual is zero, either all processes driving gait were captured or noise obscured the disambiguation of missing processes (a limit imposed by sensor technology).

### 2.4. Data quantity

To assess the data quantity required to encode *Exo* gait, we evaluated the effect of training data length on variance accounted for. The required data quantity will dictate the settings in which discrepancy modeling is practical, such as in gait analysis where datasets contain only a few gait cycles (Kerkum et al., 2015; Ries et al., 2014). Therefore, we iteratively changed training data length (*n* = [1, 10, 20, 30, 45, 60, 90, 120, 180] seconds) for the *Nominal, Exo*, and *Augmented* models and calculated *Exo* variance accounted for. The full training set contained 210 seconds of data for each participant. For all training set lengths, we evaluated models using the full 30-second validation dataset. For all variables and across individuals, we compared (i) the *Augmented* versus *Exo* models at each training data length, and (ii) the *Exo* models with the smaller versus full training dataset at each length.

## 3. Results

Discrepancy modeling captured the complex changes in gait underlying individual responses to ankle exoskeletons. Across participants, the *Nominal* model accounted for 96.0 – 98.1% and 69.7 – 76.8% of the *Nominal* condition’s kinematic and EMG median variance, respectively, while only accounting for 86.3 – 94.0% and 51.6 – 66.4% of the *Exo* condition’s kinematic and EMG median variance, suggesting the processes underlying *Nominal* and *Exo* gait differ. While joint kinematics during *Exo* gait were well predicted using the *Nominal* model (orange in Fig. 2; median *R*^2^ = 0.863 – 0.939), the *Augmented* model (green in Fig. 2) significantly increased variance accounted for (*p* < 0.042, median *R*^2^ = 0.928–0.963). For EMG, the *Augmented* model (median *R*^2^ = 0.665–0.788) accounted for significantly more variance than the *Nominal* model (median *R*^2^ = 0.516 – 0.664). Indeed, the *Augmented* model accounted for 2.34 – 14.91% more variance (median across participants) in *Exo* gait than the *Nominal* model, suggesting that the discrepancy model captured processes driving responses. The *Augmented* model also predicted kinematics and EMG with similar accuracy to the upper-limit *Exo* model (gray in Fig. 2). While the *Augmented* model explained significantly less variance in the knee, ankle, HAM, and VAS compared to the *Exo* model (Sidak *α* = 0.013), differences in median *R*^2^ were small (knee = 0.0144, ankle = 0.0267, HAM = 0.0469, VAS = 0.0936). The remaining variables’ variance was not significantly different between the *Augmented* and *Exo* models. Minimal kinematic variance was left unexplained by the *Exo* model (median *R*^2^ = 0.954 – 0.978), but only accounted for 72.4% – 81.5% of the median variance in EMG during *Exo* gait across all individuals.

**Figure 2:**
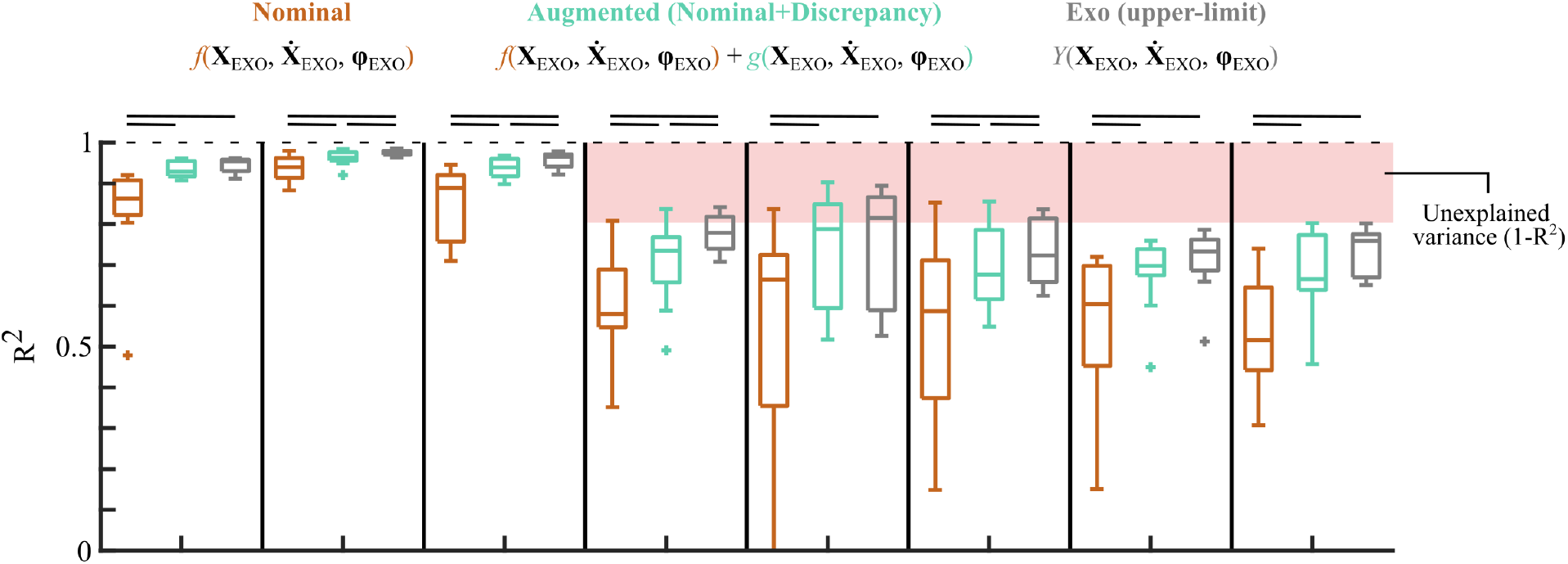
Discrepancy modeling captured individual responses to ankle exoskeletons and quantified sufficiency of kinematic and EMG measurements for encoding exoskeleton gait. The Augmented (Nominal+Discrepancy) model (green) explained significantly more variance in joint kinematics and EMG measurements compared to the Nominal model (orange). Boxplots show variance accounted for (*R*^2^) across participants (N = 11) on 30 seconds of held-out exoskeleton gait data. Horizontal bars denote significant differences between the models according to Wilcoxon Signed-Rank tests with Holm-Sidak Stepdown corrections for multiple comparisons (*α* = 0.05). The *Exo* model (gray) indicates expected maximum variance.

Residual analysis revealed non-zero trends in the stride-averaged residuals from the *Nominal* model across variables; Fig. 3 shows results from one representative subject. If the data were sufficient to fully encode the processes governing exoskeleton responses, the stride-averaged residuals would be at or near zero. Compared to the *Nominal* model (orange in Fig. 3), both the *Augmented* model (green in Fig. 3) and *Exo* (gray in Fig. 3) reduced the variance in the stride-averaged residuals, especially for the kinematic variables. For example, the mean of the *Nominal* model’s hip residual has a clear non-zero oscillatory pattern, indicative of an incomplete model. The magnitude of this pattern was reduced by 52.8% when the *Augmented* model predicted *Exo* gait. When evaluating residuals in predicted EMG, we focus on the portions of the gait cycle in which muscles are active. We observed a decrease in magnitude of the stride-averaged residuals across all muscles with the *Augmented* model.

**Figure 3:**
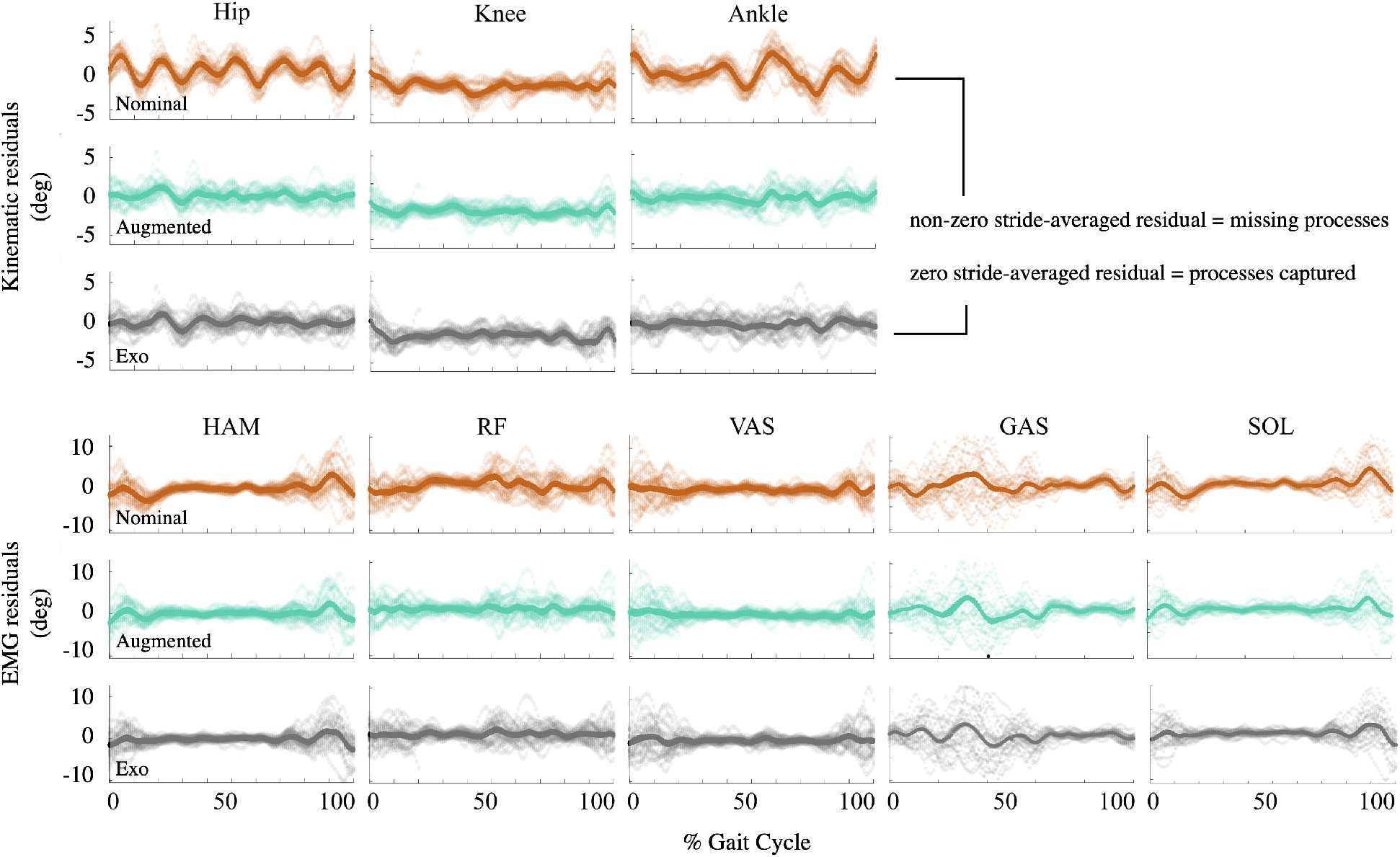
Residual analysis for one representative participant indicates discrepancy modeling identified processes underlying complex gait changes. Less saturated colors represent residual between model outputs and measured exoskeleton gait; more saturated colors represent the stride-averaged residuals. Patterns in the Augmented model’s residual diminished, as compared to Nominal, suggesting discrepancy modeling captured processes underlying gait responses to ankle exoskeletons.

Except for 1 second of training data – which predicted negative *R*^2^ for all models (Fig. 4) – the *Augmented* model did not predict significantly different *R*^2^ than the *Exo* model regardless of data quantity, for all variables and across individuals. This suggests discrepancy modeling is effective, even with small data quantities. However, the ability to encode the response with the discrepancy diminishes with less data. For all variables and across all individuals, 180 seconds of data are needed for the *Exo* model to predict statistically similar *R*^2^ to the *Exo* model’s full training dataset.

**Figure 4:**
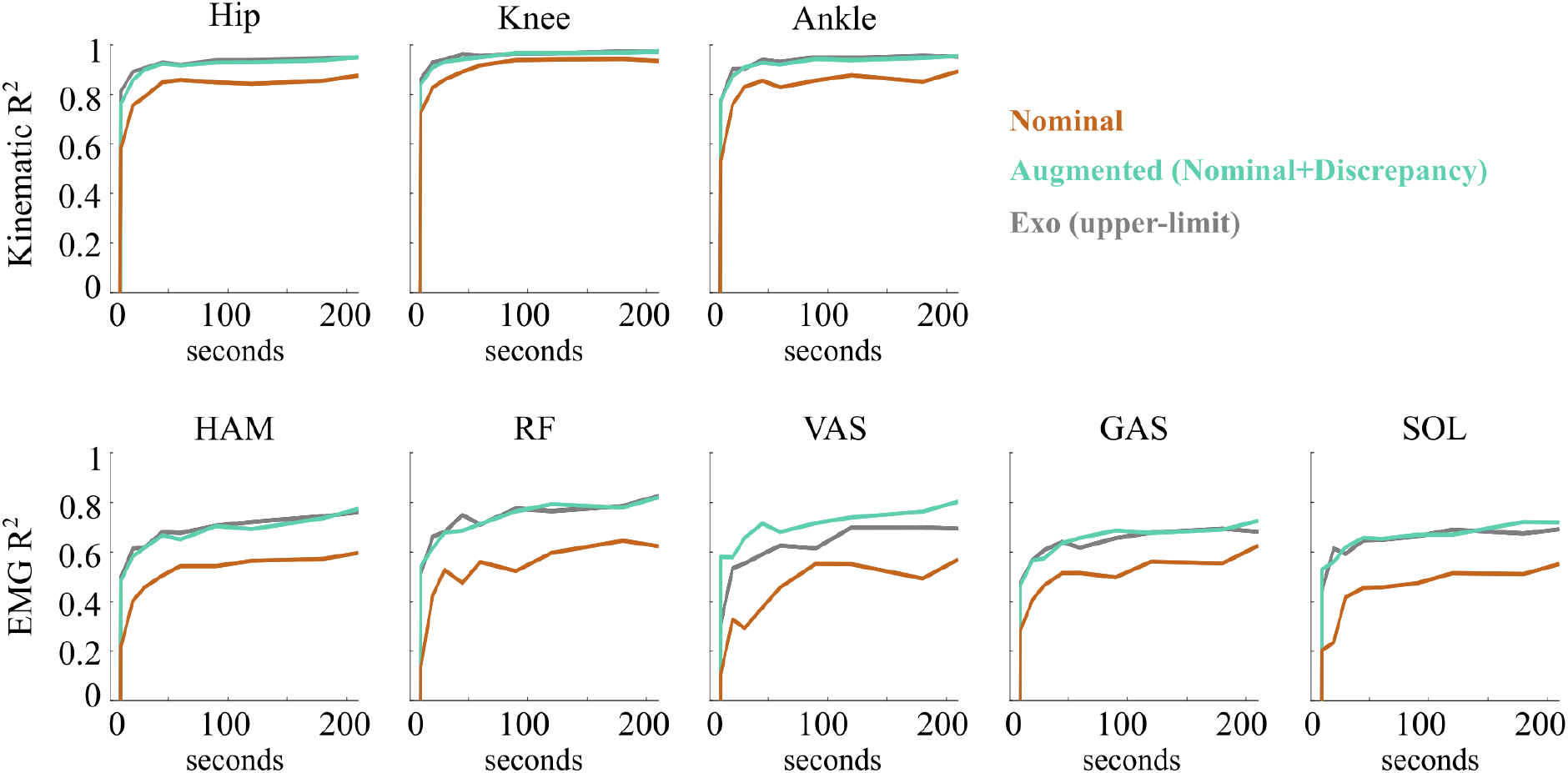
Discrepancy modeling captures exoskeleton responses, even with small training datasets. Median *R*^2^ for kinematics (top) and EMG (bottom) during exoskeleton walking for the Nominal (orange), Augmented (Nominal+Discrepancy) (green), and upper-limit Exo (gray) models over training set sizes ranging from 1 to 210 (full set) seconds.

## 4. Discussion

A neural network-based discrepancy modeling framework successfully captured changes in the processes governing gait kinematics and EMG with ankle exoskeletons (*i.e.*, discrepancies) without requiring prior knowledge or assumptions about the physiological structure of the discrepancy. Compared to the *Nominal* model alone, improved predictions of gait using with the *Augmented* (Nominal+Discrepancy) model highlights this framework’s ability to quantify exoskeleton responses as discrepancies. However, non-zero residuals, even for the upper-limit *Exo* model, indicate additional measurement modalities and/or improved measurement resolution are needed to capture the complex muscle-level changes in gait with ankle exoskeletons. Our framework’s ability to isolate discrepancies in *Exo* walking provides a first step towards leveraging machine learning to uncover physiological mechanisms driving responses to exoskeletons or other assistive devices.

As suggested by our prior work developing discrepancy models in synthetic systems (Ebers et al., 2022), we found that neural networks captured individual-specific processes governing changes in kinematics and EMG with exoskeletons from experimental data. As described below, the ability to quantify discrepancies is relevant to existing modeling frameworks because it can (i) capture differences in exoskeleton responses, (ii) encode physiological mechanisms underlying responses, and (iii) resolve dynamical inconsistencies between observations and predictions.

Our results support that discrepancy modeling can quantify individual-specific exoskeleton responses without assuming neuromusculoskeletal structure. Because few studies report kinematic and EMG accuracy, and because we predict accelerations and second time-derivatives, head-to-head comparisons are challenging. Regardless, we expect our prediction accuracy is comparable to predictions from physiological-detailed models (Falisse et al., 2019). The ability of our framework to encode discrepancies from data is relevant for modeling exoskeleton responses in patient populations: existing physiologically-detailed models are challenging to tune to an individual’s physiology and motor control, especially for individuals with highly-heterogeneous physiology (Pitto et al., 2019; Veerkamp et al., 2019; Meyer et al., 2017; Sauder et al., 2019; Nikoo and Uchida, 2022; Falisse et al., 2019; Handsfield et al., 2016). Our discrepancy framework’s ability to encode processes governing gait changes with exoskeletons could, therefore, enable precise individual-specific representations of exoskeleton walking, even in the absence of explicit understanding of an individual’s physiology. These structures may enable more accurate predictions of exoskeleton responses and suggest physiological features of gait explaining these responses (Rosenberg et al., 2020; Falisse et al., 2019; Jackson et al., 2017).

Discrepancies may encode physiological mechanisms underlying exoskeleton responses. Increased residuals when predicting *Exo* gait with the *Nominal* model suggests missing processes. Reductions in these residuals with the *Augmented* model further supports that discrepancies captured exoskeleton responses. Yet, we do not know what physiological mechanisms constitute the discrepancy. Prior studies have found motor, muscle-tendon, and optimal control mechanisms that impact responses to exoskeletons and other assistive devices (Jackson et al., 2017; Sawicki and Khan, 2015; Falisse et al., 2019). Therefore, it is likely that muscle-tendon properties, sensory processes, and motor control underlie discrepancies (Sawicki and Khan, 2015; Zajac, 1989; Geyer et al., 2006; Martelli et al., 2015; Franz, 2021; Falisse et al., 2019). However, these studies do not reveal which feature(s) underlying responses. Discrepancy models, by definition, isolate the exoskeleton response, such that the features underlying responses could be grounded in physiology. This work shows the discrepancy modeling framework can capture individual-specific processes driving changes in exoskeleton kinematics and EMG. Uncovering which of the above physiological mechanisms – and possibly others (Ting and Chiel, 2017; Schroeder et al., 2014; Schofield et al., 2020) – explain discrepancies is an interesting avenue of future research.

This discrepancy modeling framework represents a complementary approach to established biomechanical models. Existing modeling approaches typically use iterative optimization to select model parameters(*e.g.*, cost functions, muscle-tendon parameters, joint moments, and activation dynamics) explaining changes in kinematics, kinetics, and muscle activity across experimental conditions (Falisse et al., 2019; Meyer et al., 2017; Pitto et al., 2019; Rosenberg et al., 2022). Conversely, the fitting process of the discrepancy modeling framework tunes model parameters based solely on the consistency of model predictions to kinematic accelerations and EMG second time-derivatives. As a result, discrepancy modeling does not require iterative optimization: the neural networks used here could be replaced with a least-squares regression model to more-rapidly (but possibly less-accurately) discover discrepancies. Further, discrepancy modeling aims to resolve dynamical inconsistences, not just reproduce gait kinematics. Resolving dynamical inconsistencies with exoskeletons in this study is analogous to OpenSim’s residual reduction algorithm (RRA), which ensures dynamic (*i.e.*, forces and moments) consistency between a skeletal model and measured external forces (Delp et al., 2007; Koller et al., 2015; Sturdy et al., 2022). Therefore, while this work supports the validity of discrepancy modeling for exoskeleton responses, the framework’s utility may generalize to myriad biomechanical applications.

The models in this work, similar to prior work (Rosenberg et al., 2020; Pitto et al., 2019; Sauder et al., 2019), struggled to capture the complex changes in EMG. If gait with exoskeletons is not fully described by the data, it is unlikely the discrepancy will completely encode exoskeleton responses. The limited prediction accuracy of the *Exo* model (*R*^2^ < 1) indicates that kinematic and EMG data lack information needed to encode processes driving EMG. Therefore, discrepancies describing EMG responses between *Nominal* and *Exo* gait could not be comprehensively captured. Why did we not capture EMG responses? If the *Exo* model provided the expected maximum variance, and variance remained unexplained, we may need additional measurement modalities and/or improved resolution. For example, Sawicki and colleagues (2015) demonstrated that including muscle-tendon mechanics could improve predictions of individual responses to ankle exoskeletons during hopping (Sawicki and Khan, 2015). Incorporating additional data, such as measures of muscle-tendon mechanics from ultrasound (Nuckols et al., 2020) or musculoskeletal simulation could improve response predictions. The training data quantity may also limit our ability to encode from data the processes driving gait and exoskeleton responses. Particularly for EMG, unexplained *Exo* variance remained. Prediction accuracies, however, plateaued for all variables, similar to a prior study of gait with exoskeletons (Rosenberg et al., 2020), suggesting that predictions would not improve with more training data alone. Finally, experimental protocol may impact our ability to encode exoskeleton responses from data. For example, the may not have elicited large or diverse enough perturbations to distinguish processes underlying responses from noise during model fitting. Experimental paradigms eliciting larger responses may increase the ability to encode discrepancies from data.

One limitation of this study is our evaluation of a single passive exoskeleton condition, similar to current standard practice for clinical prescription of ankle foot orthoses. However, understanding whether discrepancy modeling can encode responses to changes in other exoskeleton parameters (*e.g.*, other mechanical properties or powered exoskeleton control policies) represents important areas of future work. Additionally, we applied discrepancy modeling to nondisabled participants; the framework’s ability and/or the data needed to capture discrepancies may change for individuals with disabilities (Wren et al., 2011, 2020). Further, the participant omitted from analysis highlights an important limitation of discrepancy modeling: the discrepancy relies on training data emerging from the same processes that govern observed behavior (the test data, in this study). This participant may have fatigued or adapted during the trial, such that training the discrepancy on data early in the trial may fail to capture discrepancies later in the trial. Another potential limitation was in our selection of processing parameters (*e.g.* low-pass filtering cutoff frequency), as such parameters may remove non-noise information from the signals (Wimalasena et al., 2022). Finally, while the discrepancy modeling framework is model agnostic, it is important to consider limitations of the model’s structure (*e.g.*, neural networks), which may impact the ability to capture and interpret exoskeleton responses. Therefore, we hyperparameter tuned layer quantity, nodes per layer, and activation functions to approximate gait without overfitting. Despite the network’s simple architecture, it captured discrepancies; more sophisticated deep learning architectures could be employed in future studies.

## 5. Conclusion

This study used a neural network-based discrepancy modeling framework to quantify individual responses to ankle exoskeletons. We found that discrepancy modeling captured processes underlying complex changes in kinematic and EMG data with exoskeletons. Additionally, we demonstrated how discrepancy modeling can be used to identify what and how much data are needed to capture responses. Discrepancy modeling is a unique and innovative tool that complements current biomechanical modeling approaches and may accelerate the discovery of individual-specific mechanisms driving responses to exoskeletons, other assistive devices, and clinical interventions.

## Acknowledgements

MRE acknowledges support from the National Science Foundation under award GRFP DGE-1762114. MCR acknowledges support from the National Science Foundation Graduate Research Fellowship Program under grant no. DGE-1762114 and the National Institute of Child Health and Human Development under award number F32HD108927. JNK acknowledges funding from the National Science Foundation AI Institute in Dynamic Systems grant number 2112085.

